# Finite element analysis of a neural implant for cytostatic hypothermia and a novel heat management system

**DOI:** 10.1101/2024.05.13.594046

**Authors:** Syed Faaiz Enam, Reed Chen, Ravi Bellamkonda

## Abstract

The treatment of glioblastoma (GBM) presents significant challenges, with median survival rates remaining low despite standard-of-care therapies. This study expands upon the findings of a novel approach to managing GBM, namely cytostatic hypothermia, through the computational evaluation of a fully implantable system. Our proposed system utilizes a multi-probe array and a novel artificial internal circulation system (AICS) to achieve homogeneous cooling within the brain without overheating any portion of the body. Finite-element modeling was employed to simulate bioheat transfer and fluid dynamics. Our results indicate that the multi-probe array can attain local tissue temperatures within the cytostatic range (20 to 28degC) while minimizing thermal gradients. The use of multiple narrow, thermally conductive probes enhances cooling uniformity with minimal tissue displacement. The revolutionary AICS provides a form of heat management that has not previously been attempted to the best of our knowledge. In this study, it successfully facilitates the transfer of heat from the intracranial region to the skin in the body. Future work will focus on device prototyping and validation through in vitro and in vivo studies in large animal models. These simulations suggest that the proposed intracranial cooling system makes cytostatic hypothermia a practicable approach against GBM. Furthermore, this approach to internal heat management may also open new avenues for treating neurological conditions through local and chronic hypothermia, extending beyond the short-duration (acute) cooling methods currently tested.

## 1. INTRODUCTION

The treatment of glioblastoma (GBM) remains a formidable challenge, with median survival extending to 15-18 months under standard-of-care protocols [1], [2]. The small incremental benefits offered by contemporary therapies underscore the urgent need for innovative treatment modalities. This limited success of current approaches results in virtually all GBMs recurring. However, ∼80% of recurrences occur 0-2 cm from the original site (and 50% within 0-1 cm) [3], [4], [5]. In addition, supra-total resection, i.e. resecting beyond the tumor bulk mass and into brain, results in improved survival [6], [7]. However, resecting healthy brain is not an ideal approach as it can result in functional deficits and morbidity [8]. Regardless, that recurrence is close to the resection margin highlights the potential for *local therapeutic interventions* against recurrent GBM. Our prior work has elucidated the efficacy of local *cytostatic* hypothermia in prolonging survival in two rat GBM models [9]. We and others have identified that various tumor types exhibit cell division cessation at temperatures between 20–28°C [9], [10], [11], [12], [13], [14], [15], [16]. This approach not only demonstrates promising preclinical results but also holds translational potential, as the physical principles that undergird the strategy apply similarly to rodents and to humans. Nonetheless, a pivotal challenge lies in the design of a clinically relevant medical device [9], [17], [18], [19], [20], [21].

The development of such a medical device necessitates consideration of several design parameters. Foremost, the necessity for a *fully implantable* device is underscored by the susceptibility of GBM patients to infections (rendering external components a hazard). Furthermore, the device must ensure *homogenous cooling* within the targeted tissue volume; that is, avoiding sharp temperature gradients and regions facilitating tumor proliferation (from >28°C) [9] or impair neural functionality (from <20°C) [18]. A third requirement is compatibility with magnetic resonance imaging (MRI) given the reliance on MRI for regular monitoring of tumor progression in GBM patients. Finally, the device must be compatible inasmuch that material choice and pumping of heat does not damage tissue.

Although current methodologies to induce hypothermia as a treatment are in existence, they fall short of meeting these criteria. Conventional systems involve an interface with the tissue, heat transfer through a medium to an external component, and predominantly utilize air-based convection for heat extraction [18], [19], [22], [23], [24], [25]. These systems incorporate percutaneous components and external elements, posing substantial infection risks.

In this computational study, we propose a novel system designed for the intracranial and geometrically homogenous administration of cytostatic hypothermia. This incorporates platform technology to enable heat management. Our approach uses finite-element modelling to execute simulations of heat transfer and fluid flow. To achieve a fully implantable system, we innovatively and safely employ the scalp and skin as a natural heatsink. The designs conceived are not only MR-compatible but also suggest evidence of safety regarding heat dissipation. Our results suggest that one of the major technical impediments to the clinical application of cytostatic hypothermia is surmountable.

## 1. METHODS

### a. Computational analysis

Finite-element modelling (FEM) was performed on COMSOL 5.2 under a Floating Network License at Duke University. Modules used included the Heat Transfer module and Pipe Flow module. Bioheat transfer was evaluated using Pennes’ equation within the geometric region considered to be brain or skin tissue. Heat transfer was evaluated in all non-biological materials including metal components and fluid. All tissues and materials used in the model were assumed to be homogeneous and isotropic – features such as brain blood vessels were not modeled. Multiple parameters used for the tissues, perfusion, and materials were investigated [26], [27]. Original perfusion values in units of mL/100□g/min were converted into s-1 using a blood density of 1050 kg/m3. The blood perfusion and metabolic heat generation parameters were temperature-dependent based on the following equations:

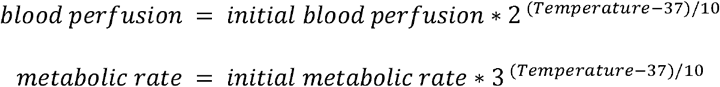

### b. Brain and array simulations

The geometry of the brain was provided by Dr. Paolo Maccarini at Duke University via segmentation of an MRI scan. The Region of Interest (ROI) was determined from literature to be 3 cm in diameter. An additional 2 cm region around this was included as a Region of Analysis wherein the mesh for analysis was finer than in other portions of the brain. The multiprobe array was modelled with a hexagonal pattern, and the size of the array, the number of probes, the pitch between the probes, the diameter of each probe, and the length of the probes were parameterized and swept over. The length of the probes would cut off once it reached the boundaries of the ROI. For an optimized model, the probes at the periphery of the hexagonal array were manually adjusted to improve cooling. The Peltier was defined by a geometry wherein a heat flux was applied so as to pull heat from the multiprobe array. BioHeat and Heat Transfer modules were used for this simulation.

### c. Passive and active heat management simulations

The geometry of the scalp and skin were assumed as flat rectangles. A Peltier was embedded in the scalp such that a heat flux was produced from the Peltier to a Waterblock. The Waterblocks were parametrically designed to consider size and number of turns of internal piping. A pipe between the two Waterblocks was parametrically designed for length. The thickness of the skin and subcutaneous adipose tissue was fixed to be 10 mm [28], [29]. Heat Transfer was used to model the transfer of heat from the Peltier to the Waterblocks and between the Waterblocks and the fluid. Bioheat Transfer was used to model the transfer of heat from the Waterblocks and tubing to the skin. The Pipe Flow module was used to model the laminar flow of fluid in the tubing.

### d. Design of full system

An envisioned system was modeled in Autodesk Fusion360 with our modelled brain and freely available models of a human torso and skull.

## 3. RESULTS

### a. Setting up the computational model

After surgical resection and radiation therapy, 80% of GBMs recur within 2 cm from the original site and 50% recur within 1 cm of the original site [3], [4], [5]. Using this knowledge and the 3-dimensional brain model, we set up geometries representing a 3-cm diameter spherical region of interest (ROI) inside the brain (**Fig. 1a and b**). This ROI is the region where GBM is likely to recur between 50 and 80% of the time. In patients after surgical resection, there may also be a variable surgical cavity at the center of the ROI. However, as this would have mostly static CSF fluid it would not affect the ability to cool significantly. In addition, due to its variability in each patient, it was not modelled. We set up a mesh to have more precision in the ROI (**Fig. 1c**). To study the dynamics more accurately around this ROI, we established a 2-cm ‘Region of Analysis’ (ROA) with increased mesh density as well (**Fig. 1c and d**). Heat transfer was modeled using the Pennes’ bioheat equation. These parameters provided a simplified representation of biological heat transfer in brain with an ROI (**Fig. 1d**).

**Figure 1.**
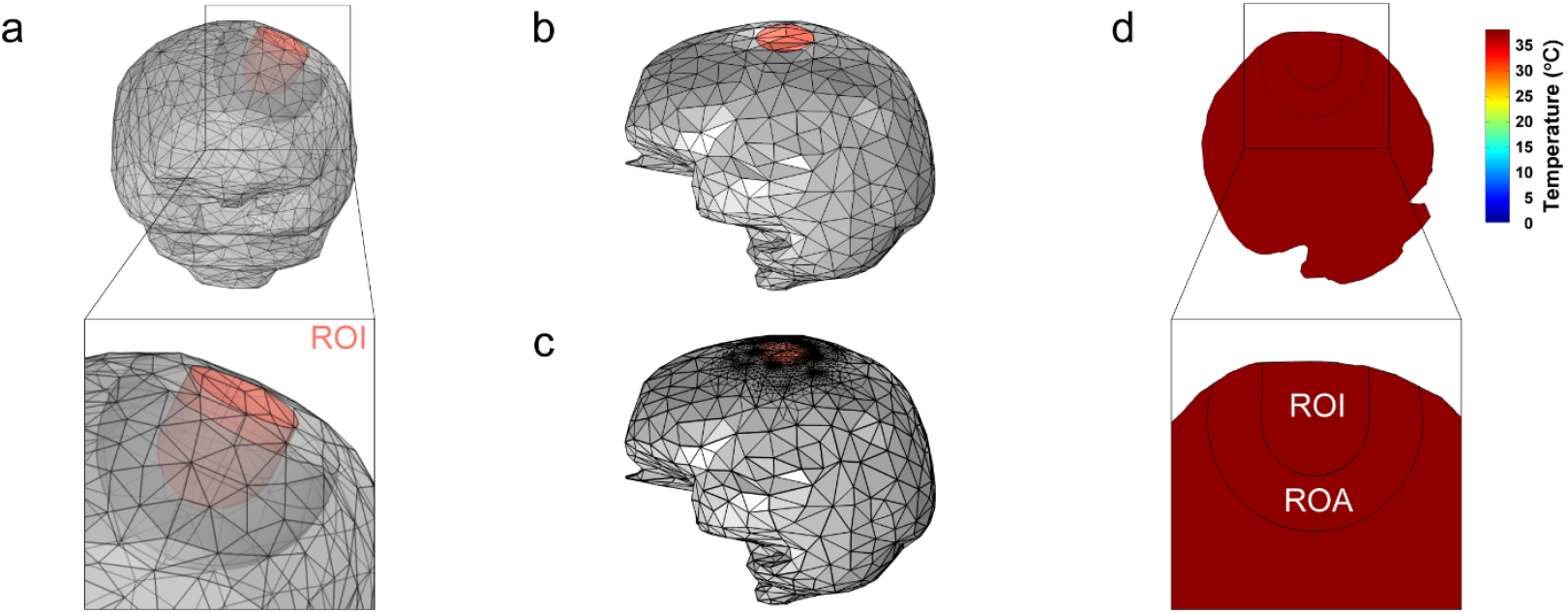
Computational model of bioheat transfer in the brain. (a) Anterior view of 3D model with geometric faces. The 3-cm diameter region-of-interest (ROI) is visible in orange. A further region outside of the ROI, the region-of-analysis (ROA) is visible. (b) Lateral view of 3D brain model. Orange ROI visible with circle around it representing ROA. (c) Mesh geometry for simulation. Increased mesh density visible in ROI and ROA. (d) Resulting coronal slice of brain after bioheat simulation, rotated to verticalize the ROI axis. Slice shows both ROI (where probes would insert) and ROA. The brain model reaches blood temperature.

### b. Modeling local cooling of tumor-infiltrated brain with a single probe

To model local intracranial cooling, we added a geometry representing a probe implanted into the ROI (**Fig. 2a**). This probe was connected to a copper base on top, outside the brain (**Fig. 2a**). The upper surface of the base incorporated a heat energy transfer boundary (ranging from 5 – 15W) to pull heat from the base and hence cool the probe. This boundary represents the boundary between the base and the cold side of a thermoelectric module (TEM or Peltier plate). A mesh was created that focused on relevant regions including the ROI and ROA (**Fig. 2b**). For the evaluation of this system, we varied the radius and thermal conductivity of the intracranial probe and heat flux across the base (**Fig. 2c**). Our target was to achieve a maximum tissue temperature of 25°C in the ROI. The probe was assumed to either be a solid metal (with thermal conductivity close to that of copper) or a heat pipe (with an estimated thermal conductivity of 10x copper).

**Figure 2.**
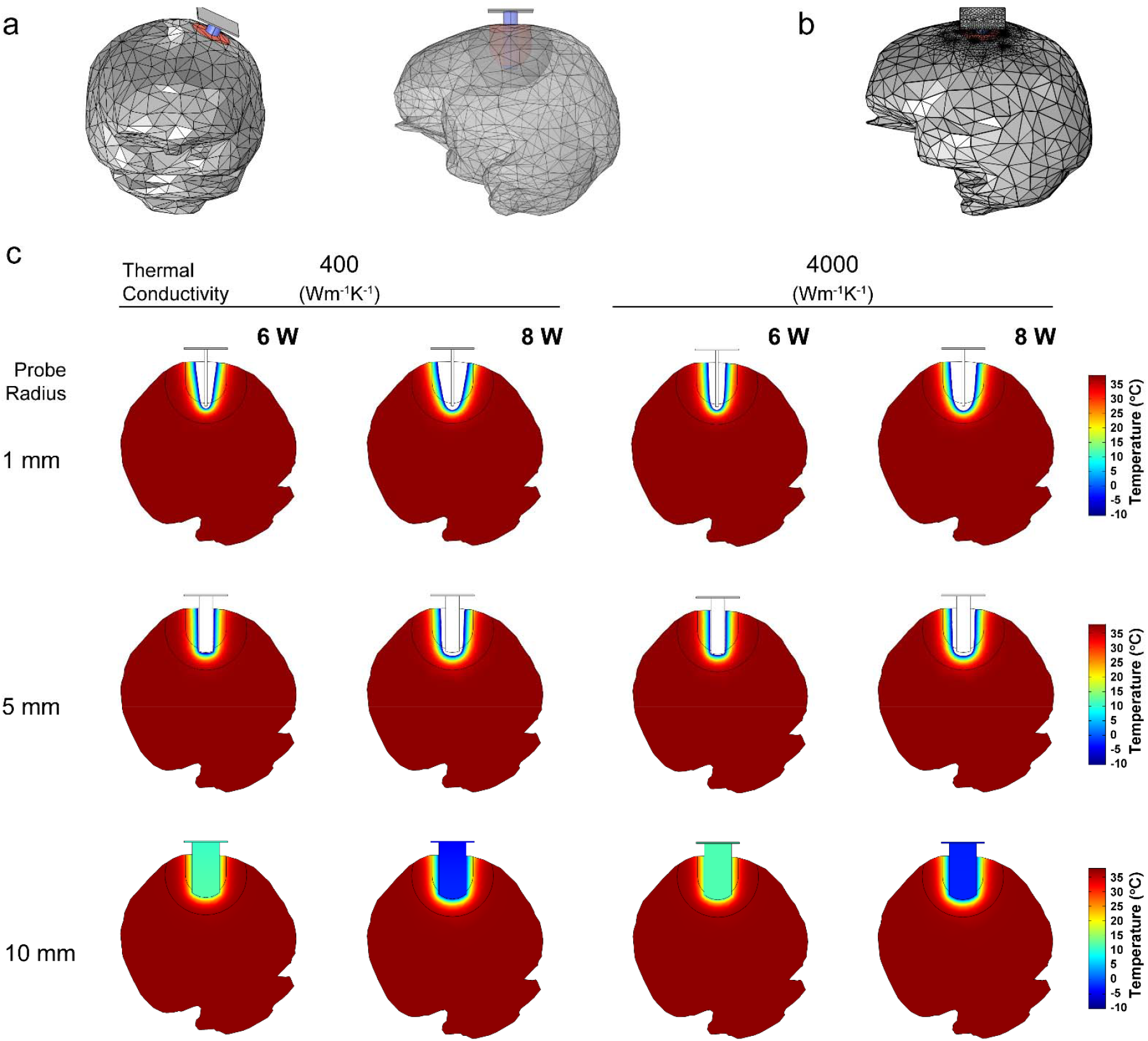
Intracranial cooling with a single probe. (a) Anterior and lateral view of 3D model with single intracranial probe (blue) and ROI (orange) and ROA is visible through the translucent lateral view. (b) Physics-informed mesh showing increased granularity around the cooling region. (c) Coronal brain slice of temperature profile with varying probe radius, thermal conductivity of the probe, and heat flux. White color indicates temperatures below the range of the scale.

The use of material with higher thermal conductivity facilitated the ability to make the cooling more uniform down the probe. This is particularly exaggerated with the cooling profile of the 1-mm radius probe (**Fig. 2c**). With a heat flux of 6 to 8W from the TEM, temperatures within a 1-mm and 5-mm radius probe and local tissue reached <0°C. This also established a very steep temperature gradient. A 10-mm radius probe was required to keep local temperatures above freezing (cytotoxic) temperatures but within cytostatic temperatures (**Fig. 2c**). While a 1-mm radius probe took up only 13 mm^3 (0.013 mL, 0.07% of ROI), the 10-mm radius probe 8738 mm^3 (8.7 mL, 50.12% of ROI) (**Supplementary Table 1**). However, the larger probe at 6W kept ROI temperatures between 11 and 29.1°C and 7W kept it between 4.3 and 26.1°C. Within this, however, only 32% of the ROI was in 20-25°C range at 6 or 7W (**Supplementary Table 1**). These data suggest that a single-probe approach is impractical due to the amount of volume it displaces and the tight therapeutic window it provides.

### c. Modeling a multi-probe strategy to improve cooling homogeneity

To prevent local subzero temperatures, improve the homogeneity of cooling, and reduce the volume of tissue displaced, we incorporated and assessed a multi-probe strategy in the ROI with a surrounding ROA (**Fig. 3a**). Hexagonal arrays of probes were investigated by modeling probes with radii of 0.0625, 0.125, 0.25, and 0.5 mm. The pitch between probes were kept consistent within an array, and across groups ranged from 1.5x to 20x the probe diameter. Heat flux between 6W and 10W was evaluated (**Supplementary Tables 2 and 3**). A physics-informed mesh was created to be more precise around interacting geometries (**Fig. 3b**). Each probe extended 2 mm past the region of interest to ensure sufficient cooling of the recurrence volume (**Fig. 3c**). With these parameters, the number of probes in each array and their diameter could be varied. Additionally, as previously, thermal conductivity of 400 W/(m*K) (**Supplementary Table 2**) and 4000 W/(m*K) (**Supplementary Table 3**) was tested. Minimum and maximum temperatures within the ROI were evaluated and displacement volume by the probes was studied.

**Figure 3.**
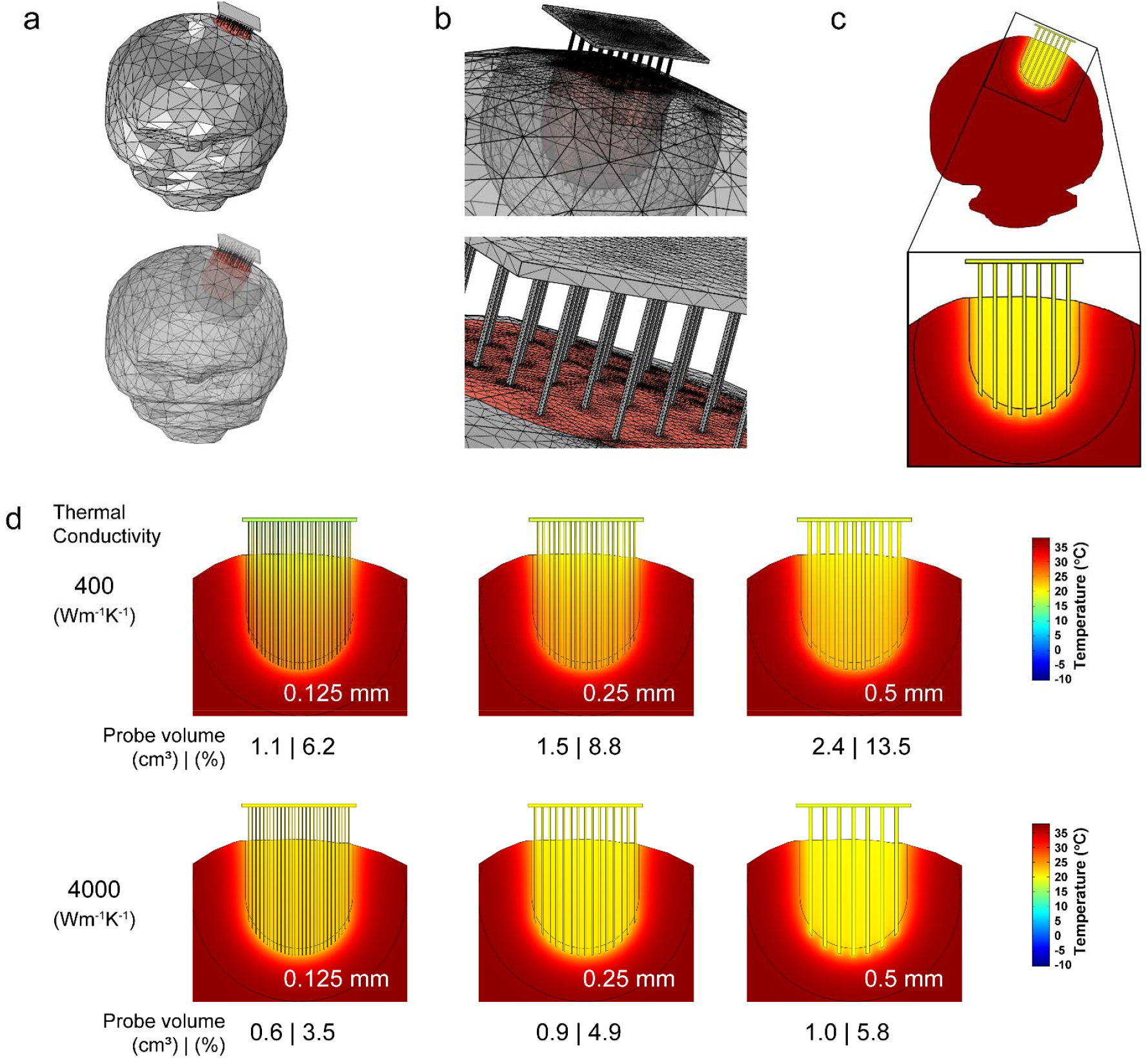
Multiprobe array for homogenous intracranial cooling. (a) Anterior view of brain with multiprobe array inserted into ROI (orange). (b) Zoomed view of interface between multiprobe array and ROI in brain. (c) Example coronal brain slice with temperature profile and inset depicting the multiprobe array. (d) Temperature profile of brain slices with multiprobe array demonstrating homogenous cooling. Across the subfigures, thermal conductivity and probe radius have been varied. Below each slice is the amount of volume that the probes take up/displace.

Results demonstrate that 6 to 8W can be sufficient to homogenously cool the ROI to be within cytostatic temperatures but above freezing temperatures (**Fig. 3d**). With probes of thermal conductivity of 400 W/(m*K), the most homogenous result, and within the cytostatic range, was an array made of 0.5-mm radii probes with 1.5 mm pitch: nearly 100% of the ROI was within 20-25°C, with a max temperature 25.4°C and min temperature of 21.0°C (**Supplementary Table 2**). However, to achieve this required a total probe volume of 6735 mm^3 (6.7 mL, 38.20% of ROI). Reducing the total probe volume to ∼3.5 mL or ∼20% or ROI (possible with 0.5-mm with 2 mm pitch or 0.25 mm radius with 1 mm pitch) kept 93 to 91% of ROI within 20-25°C. In these examples, the remaining volume of tissue was <27.4°C which may still be cytostatic [12]. Smaller volume displacement was achievable if we increase the allowance of our therapeutic window to be between 15 and 30°C. For example, 0.125-mm radii probes with 0.75 mm pitch at 7W would take 1099mm^3 (1.1 mL or 6.23% of ROI) with 72% of ROI within 20-25°C (**Fig. 3d**). The remaining ROI was above 19.1°C and below 29.0°C (**Supplementary Table 2**). Alternatively, 0.0625-mm radii probes with 0.5 mm pitch at 7W took 618 mm^3 (0.6 mL, 3.5% of ROI) with 48% being between 20-25°C and the remaining between 15.5°C and 29.2°C. Probes with an increased thermal conductivity of 4000 W/(m*K), had a lower displacement volume (**Supplementary Table 3 and Fig. 3d**) to achieve similar cooling efficacy. For example, 0.0625-mm radii probes with 0.5 mm pitch at 8W would take 618mm^3 (0.6 mL, 3.5% of ROI) with nearly 100% of ROI being between 20-25°C and maximum temperature of 25.7°C (**Supplementary Table 3**). These results suggest that the use of probes with thermal conductivity of 400 W/(m*K) is feasible but will require adjustments (e.g. volume of probes or therapeutic window) while those with higher thermal conductivity readily meet ideal goals. To reasonably cool a 3-cm ROI of brain to cytostatic temperatures requires 6 to 8W of heat flux.

### d. Modeling passive and active heat dissipation to and through the skin

The prior results demonstrate that an ROI in the brain can be cooled homogenously by pulling 6 to 8W. However, this heat then accumulates in and must be removed from the TEM itself. Instead of exposing the TEM to the outside air, here we leverage the skin as a heat sink and model its ability to effectively dissipate the heat without overheating the skin.

Apart for the heat power that needs to be removed from the ROI, powering the TEM produces heat as well due to inefficiencies. Assuming a 0.5 coefficient of performance of the TEM, the additional heat flux generated by the TEM could range from 12W to 16W. Adding the heat pulled from the ROI, the total heat that needs to be dissipated the skin is 18W to 24W. Therefore, all subsequent simulations of heat transfer from the TEM were conducted assuming a heat flux of 15 – 25W. In addition, to prevent skin damage with 24 h/d of use, the maximum tolerated temperature was determined to be 40°C. This is based on equations to calculate heat damage from thermal dosage [30], [31]. We also varied environment temperatures (20, 25, 30, and 35°C) and blood perfusion of the skin (0.000333 1/s, 0.0014 1/s, and 0.010314 1/s).

To transfer heat from the TEM to the skin uniformly, we added a 1-mm thick heat sink in the form of a ‘vapor chamber’ or ‘heat spreader’ to the geometry beneath the scalp (**Fig. 4a**). As the vapor space of a vapor chamber has high thermal conductivity, an effective thermal conductivity of 5000 W/(m*K) was assumed for the vapor chamber. Heat flux was applied at the boundary of the TEM (Peltier) and the vapor chamber. On top of the scalp, a plane was designated as a potential boundary for a ‘cooling cap’. A physics-based mesh was created for this model (**Fig. 4b**). An environment with room temperature conditions (25°C) resulted in a surface skin temperature of 34 to 36°C with no power applied to the system (**Fig 4c**). The bone of the skull remained around 37°C (**Supplementary Table** 4). The bottom of the model was thermally insulated. Skin and subcutaneous adipose tissue were modeled as a 10 mm thick layer [28], [29]. The surface of the skin was cooled with a convective heat flux boundary condition. CSF was modeled with the properties of water.

**Figure 4.**
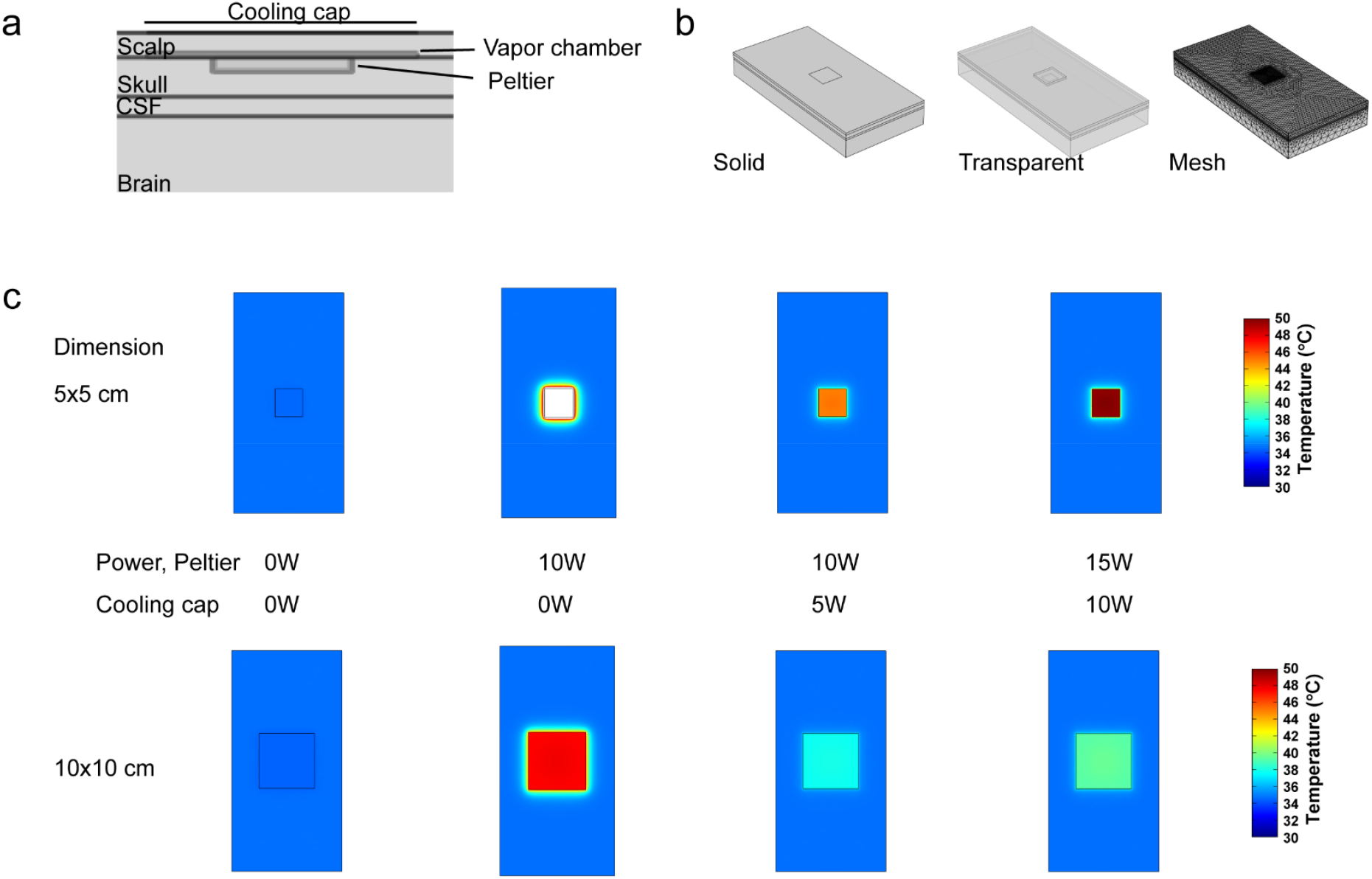
Passive cooling cap. (a) Transverse section of geometry setup. This consists of the layers in the head (scalp, skull, CSF, and brain), hot-side components of the neural device (Peltier and Vapor chamber), and a boundary region on the scalp that functions as the cooling cap. (b) Solid, transparent, and mesh representations of cooling cap setup. (c) Simulations of cooling cap efficacy in preventing overheating of the scalp. Varied are the dimension of the Peltier (and simultaneously cooling cap) and the heat flux through the Peltier and the cooling cap.

Transferring 10W of heat to the scalp raised temperatures above 50°C with a 5x5 cm vapor chamber; however a 10x10 vapor chamber also resulted in unacceptably high temperature ∼47°C (**Fig. 4c**). To counter this temperature increase, an external cooling cap (without percutaneous components and powered externally), was modelled. 10W of heat from the TEM could be safely managed with a 10x10 cm vapor chamber and an equivalently sized cooling cap pulling 5W of heat. However, higher heat transfer from the TEM requires progressively higher power from the cooling cap (**Supplementary Table 4**). Additionally, larger vapor chambers are difficult to incorporate given the space constraints of the scalp and head.

### e. Modeling an internal cooling circulation system

To improve the ability to dissipate heat safely to the skin, we looked beyond the scalp and displaced heat across a larger area of skin. We added a 5-mm thick waterblock with internal pipes located directly over the vapor chamber and beneath the scalp (**Fig. 5a**). With a 10-mm scalp thickness, 1 mm vapor chamber + 5 mm waterblock took the place of 6 mm of tissue, leaving 4 mm of scalp above the implant for this model [28], [29]. The water in this block was intended to carry heat to a second pipe-embedded waterblock located at a 160 cm distance (representing being implanted in the subcutaneous region on a patient’s dorsum). To simulate the ideal representation, one model was set such that the circuit was left ‘open’ and the input water to the waterblock had ‘returned’ to body temperature of 37°C (**Fig. 5b**). In the second, the circuit was ‘closed’ to account for the heat that would accumulate in the circulation system as a better representation of reality (**Fig. 5c**). The two waterblocks were connected with flexible tubing with an ID of 4.5mm and OD of 5mm and thermal conductivity of 0.25W/(m*K). Through this tubing, water was circulated via an implanted pump at flow velocities of 5 mL/s, 7.5 mL/s, and 10 mL/s. Combined, we term this an Artificial Internal Circulation System (AICS).

**Figure 5.**
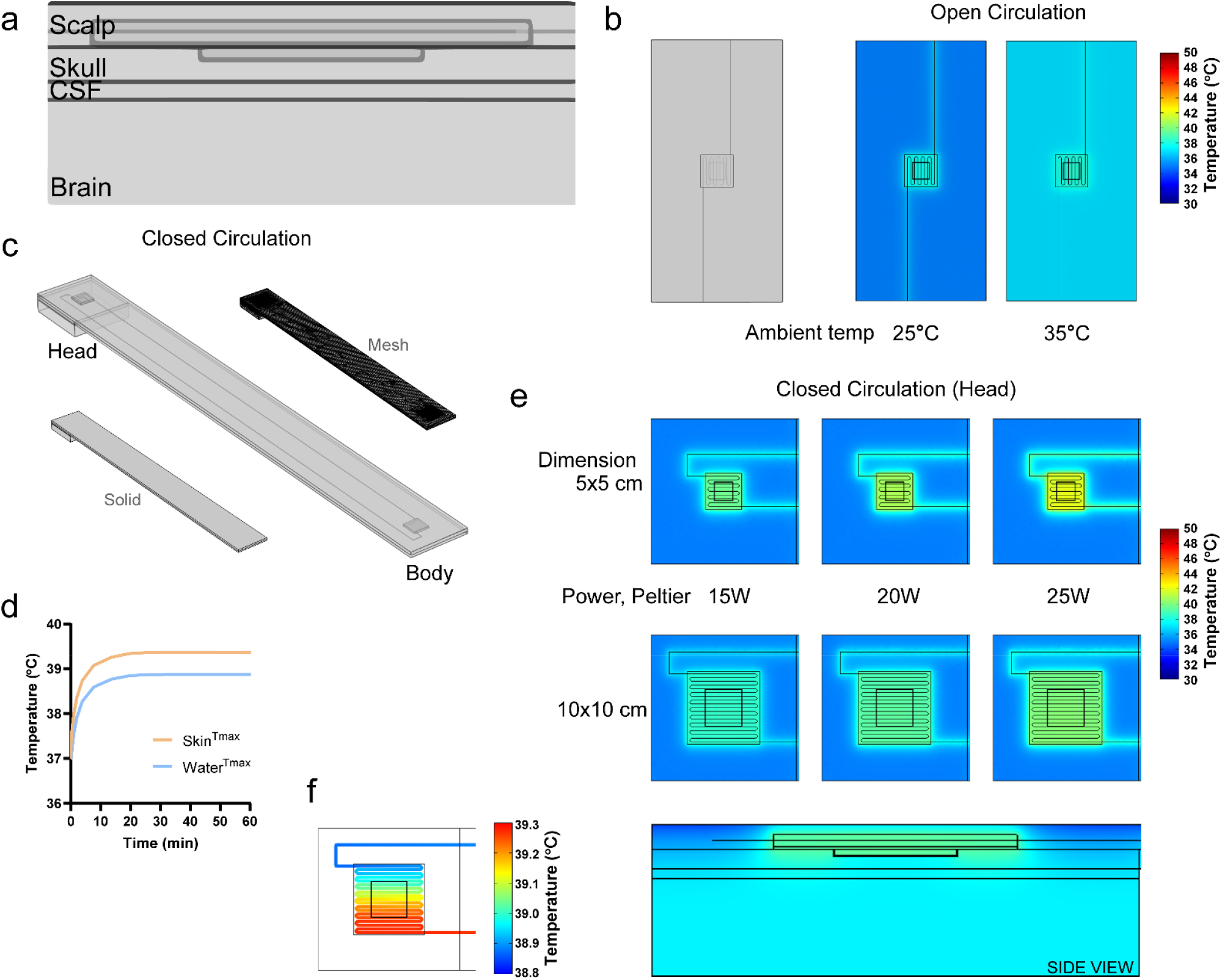
Simulations of the Artificial Internal Circulation System. (a) Transverse section of geometry setup. This consists of the layers in the head (scalp, skull, CSF, and brain), and hot-side components of the neural device (Peltier and waterblock). (b) Overhead view of geometry and simulation of scalp waterblock with an open circulation under two different ambient temperatures. (c) Solid and transparent geometry of the closed circulation system with a waterblock in the head and body connected via tubing. Physics-based mesh in depicted adjacent. (d) Graph demonstrating the time to steady-state of max temperature in skin and water. (e) Overhead and side views of simulation at the scalp waterblock with a closed circulation. Varied here are the dimensions of the waterblock and the power/heat flux from the Peltier. (f) Difference in temperature between the entrance and exit of the scalp waterblock.

**Figure 6.**
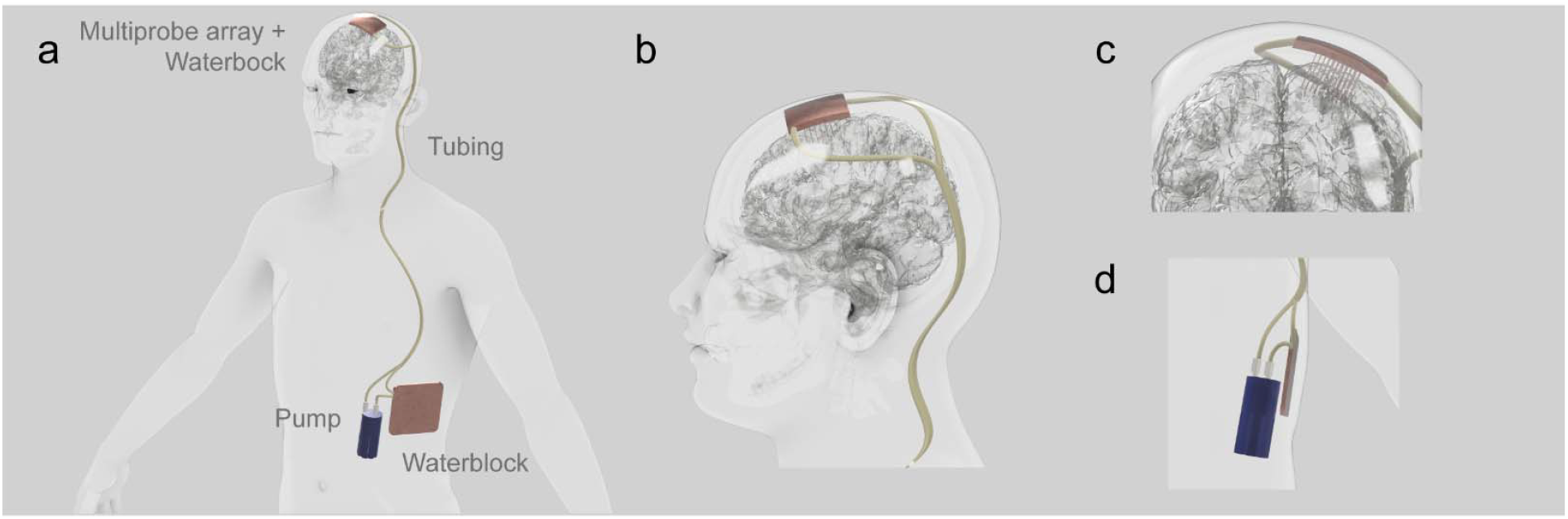
Envisioned implantable heat management system for intracranial cooling. (a) Full body view. This shows the major components of the system to enable heat transfer from the multiprobe array to the waterblock in the body. (b) Lateral view of the head. This shows how the tubing could run under the skin but outside the skull. (c) Zoomed anterior view of head with brain. Visible is the neural device with a multiprobe array and tubing. (d) Zoomed view of the pump and waterblock in the body.

In our first set of simulations, we left the tubing circuit open. This let us assume that the temperature of the fluid entering the heat sink was body temperature. We varied heat sink dimensions and fluid flow rate to be able to transfer 15 – 25W of heat without raising scalp/skin temperature > 40°C (**Supplementary Table 5**). Minimum dimensions of 5x5 cm effectively transferred up to 25W without raising temperature to >40°C at a flow rate of 7.5 mL/s at both room temperature and elevated ambient temperature of 35°C (this was also effective at 30W) (**Fig. 5b and Supplementary Table 5**). This suggests that if we can dissipate the heat generated effectively, prior to the coolant returning to the waterblock, heat transfer will be effective.

To account for incomplete heat dispersion from the circuit and better represent the AICS, we closed the fluid circuit. After closing the cooling circuit, heat accumulated in the liquid and reached steady-state in 20-30 minutes (**Fig. 5d**). At a flow rate of 10 ml/s, 15W of heat was effectively dispersed with 7.5x7.5 cm waterblocks (one at the scalp, and one on the body) (**Supplementary Table 6**). For 20W, 10x10 cm waterblocks were required to keep temperatures <40°C (**Fig. 5e and Supplementary Table 6**). The temperature drop in the coolant across an individual block was 0.5°C (**Fig. 5f**). A total of 25W of heat transfer would require a larger combination of waterblocks (**Supplementary Table 6**). This heat exchange was effective at a range of environmental temperatures. These data demonstrate that heat pulled from an ROI in the brain, along with heat generated from powering components (e.g. the TEM) can be effectively and safely dissipated into the skin of the body elsewhere.

## 4. DISCUSSION

Here we provide evidence supporting the potential of uniformly administering cytostatic hypothermia through the design of a fully implantable cooling system. Our findings underscore the benefit of employing multiple, narrow, thermally conductive probes to uniformly extract heat across substantial volumes of brain tissue. These ensure that the minimal volume is displaced while remaining in a safe and effective thermal window. Moreover, our models reveal the innovative concept of utilizing the body’s skin as a heat dissipater. This is facilitated by an artificial internal circulating system, thereby positing a novel approach to enable cytostatic hypothermia for glioblastoma treatment and future therapeutics and implants that would benefit from heat management.

Our analyses of single and multi-probe strategies elucidate the superior performance of the latter in ensuring cooling homogeneity, safe minimum temperature thresholds, effective maximum temperature thresholds, and minimizing volume displacement. If sufficiently thermally conductive, we observed enhanced outcomes in all evaluated metrics by decreasing probe diameter while increasing the quantity of probes (done by reducing pitch between probes). This is intuitively explained by an increased heat transfer surface area to volume ratio. Naturally, however, a multi-probe strategy increases the complexity of implantation (and more so when the geometry of a brain tumor can differ from patient to patient). These problems may be resolved with the development of predictive software and the precision and adaptability of advanced surgical robotic systems (e.g. such as the one developed by Neuralink). In Fig. 3d and Supplementary Table 3, the use of probes with thermal conductivity of 400 W/(m*K) frequently left a small amount of tissue at the periphery a slightly higher temperature than desirable. Some of this may still be effectively cytostatic as 28°C has been seen to stop tumor growth as well [12]. However, an alternative strategy could include increasing the span of the array so that the ROI is fully covered. The downside of this approach is the total volume displaced increases (although, without an increase in the density of the array). In the case of a GBM patient who has their tumor resected, a large cavity is frequently left behind; this lost volume may accommodate the volume displaced by the probes. Another strategy could be to reduce the pitch between the probes closer to the boundaries and hence an increased density of probes where it is required. These conclusions suggest that a multi-probe array may be effective in achieving the goals of cytostatic hypothermia and also achievable.

Of the variables that can be adjusted with the multi-probe array, thermal conductivity is perhaps the most powerful and feasible – this would be done with choice of probe material. In our simulations we used the thermal conductivity of copper; while this is the fastest conducting metal, it is also cytotoxic. To resolve this, the copper probes can be electroplated with a thin layer of palladium or platinum or coated with a bioinert material. If the layer is only microscopically thick (e.g. a few atomic layers of platinum), it should not affect the bulk thermal conductivity of the system. Heat pipes are particularly effective at heat conduction but commercially available heat pipes have diameters in the millimeter range. There exist, however, micro heat pipes, which have proven effective in electrical applications and may be applicable here [32], [33]. Alternatively, conventional solid metal electrodes could be substituted with pitch-based carbon fibers or carbon nanotubes (CNTs) with their superior thermal conductivity. These materials have been used in neural interfaces previously and could be modified to function in this application [34].

The second major output of this work is the innovative management of heat via the Artificial Internal Circulation System (AICS). Traditional methods for biological heat management often involve direct heat transfer to the outside environment via a percutaneous route – this poses significant infection risks (and more so for immunocompromised individuals). Our simulations suggest an approach that eliminates the need for percutaneous components by employing either an external cooling system, like a cooling cap, (Fig. 4) or, ideally, the AICS (Fig. 5). The deployment of an external cooling system would consist of additional TEM technology and thus require an additional power source. Additionally, it is unclear what the temperature sensation for the patient would be at the internal (in contact with a hot waterblock) and external surfaces (in contact with the cold external system) of the scalp. While pain could be abrogated via nerve block, long-term scalp damage from heating/cooling would need to be evaluated.

Alternatively, an AICS utilizes a larger surface area of the skin; by spreading heat across a larger surface, local temperatures remain at safer levels. We determined 40°C as the safety level based on thermal dosage equations [31] with the system running 24 hours per day. Based on that equation, if intermittent cooling is sufficient (as our study suggested 18h/d of hypothermia could be cytostatic), a higher temperature could be safe. Heat can be further dissipated by increasing the number or size of the waterblocks subcutaneously implanted on the body. Length of tubing can be increased (and perhaps coiled) to increase surface area contact with skin. Larger tube diameters could increase volume of water flow but this would come at the cost of patient comfort from thicker tubes and thicker waterblocks. In addition, a relatively low thermal conductivity common for flexible tubing of 0.25 W/(m*K) was used. Alternative materials for the tubing may improve thermal conductivity. Regardless, this system would be more energetically efficient than an external cooling system as it only requires circulating water with a pump. To the best of our knowledge, the AICS does not currently exist. However, it has the potential to enable not only a fully-implantable cytostatic hypothermia system for glioblastoma but also enable chronic hypothermia treatment for other diseases (e.g. injury, inflammation, or pain) and enable the development of high-powered low-noise implants.

Other considerations that may be readily solved are MR compatibility and battery power. Patients with GBM require regular MR scans to monitor tumor progression. The components of a multi-probe array and waterblocks (non-magnetic metal or carbon based) and TEM (non-magnetic semiconductor and ceramics) are MR compatible. However, the pump driving the AICS needs to be considered. Pumps classically contain a mechanical motor consisting of a permanent magnet which cannot be brought near an MRI machine. However, these could readily be substituted with non-magnetic piezoelectric pumps. Second, powering all systems may be energy intensive. The highest power consuming medical implant at this time is a Left Ventricular Assist Device (LVAD). This is powered by external rechargeable batteries and must run continuously or the heart would stop and the patient dies [35]. In the case of cytostatic hypothermia, any duration of hypothermia may prolong survival and a transient failure of the system at worst may only return the patient to status quo. Additionally, active research into higher power density and solid-state batteries is underway [36]. To avoid percutaneous power delivery, wireless power solutions already exist for the LVAD [37], [38] and could be translated here. These design considerations would make the medical device safer and more useful for physicians and patients.

There are limitations to the conclusions that can be drawn, and these could be addressed in future studies. First, we used the Pennes bioheat model to simulate heat transfer in tissue; while the perfusion component of the equation is a reasonable approximation for biological perfusion, it does not account for large vessel vasculature in the geometry. A study comparing the Pennes’ equations to a countercurrent vascular model found that while local temperatures may be different around vessels, the total power required to cool the region regardless of vessel size was the same [39]. Second, the scope of this work was limited to *in silico* simulations. Real world experimentation is required to further validate the conclusions. This would require device fabrication and the development of either an *in vitro* model of thermal brain perfusion (which would have its own limitations) or move directly to *in vivo* experiments in a large animal model (although pigs, sheep, goat, and dogs all have brain sizes smaller than humans).

In conclusion, our simulations pave the way for a novel implantable system aimed at halting glioblastoma progression through cytostatic hypothermia. The integration of a multi-probe array with an AICS holds significant promise. While our results are promising, they serve as a preliminary step, necessitating further development through prototyping and validation in large animal studies. If successful, this technology holds the potential to significantly prolong the lives of patients with glioblastoma. In addition, this approach to biological heat management may open new avenues of treatment of disease with chronic hypothermia and enable the embedding of high-power, heat generating implants.

## References

[1] M. Koshy et al., “Improved survival time trends for glioblastoma using the SEER 17 population-based registries.,” J. Neurooncol., vol. 107, no. 1, pp. 207–12, Mar. 2012, doi: 10.1007/s11060-011-0738-7.

[2] Q. T. Ostrom et al., “CBTRUS Statistical Report: Primary Brain and Other Central Nervous System Tumors Diagnosed in the United States in 2012–2016,” Neuro-Oncol., vol. 21, no. Supplement_5, pp. v1–v100, Nov. 2019, doi: 10.1093/neuonc/noz150.

[3] M. Rapp, J. Baernreuther, B. Turowski, H. J. Steiger, M. Sabel, and M. A. Kamp, “Recurrence Pattern Analysis of Primary Glioblastoma,” World Neurosurg., vol. 103, pp. 733–740, Jul. 2017, doi: 10.1016/j.wneu.2017.04.053.

[4] J. P. Kirkpatrick, N. N. Laack, H. A. Shih, and V. Gondi, “Management of GBM: a problem of local recurrence,” J. Neurooncol., vol. 134, no. 3, pp. 487–493, Sep. 2017, doi: 10.1007/s11060-016-2347-y.

[5] A. A. Brandes et al., “Recurrence pattern after temozolomide concomitant with and adjuvant to radiotherapy in newly diagnosed patients with glioblastoma: Correlation with MGMT promoter methylation status,” J. Clin. Oncol., vol. 27, no. 8, pp. 1275–1279, Mar. 2009, doi: 10.1200/JCO.2008.19.4969.

[6] J. F. Mier-García, S. Ospina-Santa, J. Orozco-Mera, R. Ma, and P. Plaha, “Supramaximal versus gross total resection in Glioblastoma, IDH wild-type and Astrocytoma, IDH-mutant, grade 4, effect on overall and progression free survival: systematic review and meta-analysis,” J. Neurooncol., vol. 164, no. 1, pp. 31–41, Aug. 2023, doi: 10.1007/s11060-023-04409-0.

[7] J. S. Young, R. A. Morshed, S. L. Hervey-Jumper, and M. S. Berger, “The surgical management of diffuse gliomas: Current state of neurosurgical management and future directions,” Neuro-Oncol., vol. 25, no. 12, pp. 2117–2133, Dec. 2023, doi: 10.1093/neuonc/noad133.

[8] P. Karschnia et al., “Prognostic evaluation of re-resection for recurrent glioblastoma using the novel RANO classification for extent of resection: A report of the RANO resect group,” Neuro-Oncol., vol. 25, no. 9, pp. 1672–1685, May 2023, doi: 10.1093/neuonc/noad074.

[9] S. F. Enam et al., “Cytostatic hypothermia and its impact on glioblastoma and survival,” Sci. Adv., vol. 8, no. 47, p. eabq4882, Nov. 2022, doi: 10.1126/sciadv.abq4882.

[10] D. Kalamida, I. V. Karagounis, A. Mitrakas, S. Kalamida, A. Giatromanolaki, and M. I. Koukourakis, “Fever-range hyperthermia vs. hypothermia effect on cancer cell viability, proliferation and HSP90 expression,” PLoS ONE, vol. 10, no. 1, p. e0116021, Jan. 2015, doi: 10.1371/journal.pone.0116021.

[11] Z. Matijasevic, “Selective protection of non-cancer cells by hypothermia,” Anticancer Res., vol. 22, no. 6 A, pp. 3267–3272, 2002.

[12] C. Fulbert, C. Gaude, E. Sulpice, S. Chabardès, and D. Ratel, “Moderate hypothermia inhibits both proliferation and migration of human glioblastoma cells,” J. Neurooncol., vol. 144, no. 3, pp. 489–499, Sep. 2019, doi: 10.1007/s11060-019-03263-3.

[13] X. Zhang et al., “Effect of mild hypothermia on breast cancer cells adhesion and migration,” Biosci. Trends, vol. 6, no. 6, pp. 313–324, 2012, doi: 10.5582/bst.2012.v6.6.313.

[14] D. K. Kelleher, C. Nauth, O. Thews, W. Krueger, and P. Vaupel, “Localized hypothermia: impact on oxygenation, microregional perfusion, metabolic and bioenergetic status of subcutaneous rat tumours.,” Br. J. Cancer, vol. 78, no. 1, pp. 56–61, 1998.

[15] C. Fulbert, S. Chabardès, and D. Ratel, “Adjuvant therapeutic potential of moderate hypothermia for glioblastoma,” J. Neurooncol., vol. 1, no. 3, pp. 1–16, Mar. 2021, doi: 10.1007/s11060-021-03704-y.

[16] T. Fay, “Early experiences with local and generalized refrigeration of the human brain,” J. Neurourgery, vol. 16, no. 3, pp. 239–260, 1959, doi: 10.3171/jns.1959.16.3.0239.

[17] G. F. Rowbotham, A. L. Haigh, and W. G. Leslie, “Cooling cannula for use in the treatment of cerebral neoplasms,” The Lancet, vol. 273, no. 7062, pp. 12–15, 1959, doi: 10.1016/S0140-6736(59)90976-6.

[18] S. G. Lomber, B. R. Payne, and J. A. Horel, “The cryoloop: An adaptable reversible cooling deactivation method for behavioral or electrophysiological assessment of neural function,” J. Neurosci. Methods, vol. 86, no. 2, pp. 179–194, 1999, doi: 10.1016/S0165-0270(98)00165-4.

[19] M. D. Smyth and S. M. Rothman, “Focal Cooling Devices for the Surgical Treatment of Epilepsy,” Neurosurg. Clin. N. Am., vol. 22, no. 4, pp. 533–546, Oct. 2011, doi: 10.1016/j.nec.2011.07.011.

[20] K. M. Karkar, P. A. Garcia, L. M. Bateman, M. D. Smyth, N. M. Barbaro, and M. Berger, “Focal Cooling Suppresses Spontaneous Epileptiform Activity without Changing the Cortical Motor Threshold,” Epilepsia, vol. 43, no. 8, pp. 932–935, Aug. 2002, doi: 10.1046/j.1528-1157.2002.03902.x.

[21] X. F. Yang, B. R. Kennedy, S. G. Lomber, R. E. Schmidt, and S. M. Rothman, “Cooling produces minimal neuropathology in neocortex and hippocampus,” Neurobiol. Dis., vol. 23, no. 3, pp. 637–643, 2006, doi: 10.1016/j.nbd.2006.05.006.

[22] D. Aronov and M. S. Fee, “Analyzing the dynamics of brain circuits with temperature: Design and implementation of a miniature thermoelectric device,” J. Neurosci. Methods, vol. 197, no. 1, pp. 32–47, 2011, doi: 10.1016/j.jneumeth.2011.01.024.

[23] D. F. Cooke et al., “Fabrication of an inexpensive, implantable cooling device for reversible brain deactivation in animals ranging from rodents to primates,” J. Neurophysiol., vol. 107, no. 12, pp. 3543–3558, Jun. 2012, doi: 10.1152/jn.01101.2011.

[24] H. Imoto et al., “Use of a Peltier chip with a newly devised local brain-cooling system for neocortical seizures in the rat. Technical note,” J Neurosurg, vol. 104, no. 1, pp. 150–156, 2006, doi: 10.3171/jns.2006.104.1.150.

[25] H. E. Bakken, H. Kawasaki, H. Oya, J. D. W. Greenlee, and M. A. Howard, “A device for cooling localized regions of human cerebral cortex,” J. Neurosurg., vol. 99, no. 3, pp. 604–608, Sep. 2003, doi: 10.3171/jns.2003.99.3.0604.

[26] A. B. C. G. Silva, L. C. Wrobel, and F. L. B. Ribeiro, “A thermoregulation model for whole body cooling hypothermia,” J. Therm. Biol., vol. 78, pp. 122–130, Dec. 2018, doi: 10.1016/j.jtherbio.2018.08.019.

[27] P. Hasgall et al., “IT’IS Database for thermal and electromagnetic parameters of biological tissues.” [Online]. Available: itis.swiss/database

[28] O. J. Ungar, U. Amit, O. Cavel, Y. Oron, and O. Handzel, “Age-dependent variations of scalp thickness in the area designated for a cochlear implant receiver stimulator,” Laryngoscope Investig. Otolaryngol., vol. 3, no. 6, pp. 496–499, 2018, doi: 10.1002/lio2.218.

[29] P. Störchle, W. Müller, M. Sengeis, S. Lackner, S. Holasek, and A. Fürhapter-Rieger, “Measurement of mean subcutaneous fat thickness: eight standardised ultrasound sites compared to 216 randomly selected sites,” Sci. Rep., vol. 8, no. 1, p. 16268, Nov. 2018, doi: 10.1038/s41598-018-34213-0.

[30] S. A. Sapareto and W. C. Dewey, “Thermal dose determination in cancer therapy,” Int. J. Radiat. Oncol., vol. 10, no. 6, pp. 787–800, Apr. 1984, doi: 10.1016/0360-3016(84)90379-1.

[31] G. C. van Rhoon, T. Samaras, P. S. Yarmolenko, M. W. Dewhirst, E. Neufeld, and N. Kuster, “CEM43°C thermal dose thresholds: a potential guide for magnetic resonance radiofrequency exposure levels?,” Eur. Radiol., vol. 23, no. 8, pp. 2215–2227, Aug. 2013, doi: 10.1007/s00330-013-2825-y.

[32] L. L. Vasiliev, “Micro and miniature heat pipes – Electronic component coolers,” Appl. Therm. Eng., vol. 28, no. 4, pp. 266–273, Mar. 2008, doi: 10.1016/j.applthermaleng.2006.02.023.

[33] M. L. Berre, S. Launay, V. Sartre, and M. Lallemand, “Fabrication and experimental investigation of silicon micro heat pipes for cooling electronics,” J. Micromechanics Microengineering, vol. 13, no. 3, p. 436, Mar. 2003, doi: 10.1088/0960-1317/13/3/313.

[34] M. Devi et al., “Carbon-based neural electrodes: promises and challenges,” J. Neural Eng., vol. 18, no. 4, p. 041007, Sep. 2021, doi: 10.1088/1741-2552/ac1e45.

[35] R. L. Kormos et al., “Left Ventricular Assist Device Malfunctions: It Is More Than Just the Pump,” Circulation, vol. 136, no. 18, pp. 1714–1725, Oct. 2017, doi: 10.1161/CIRCULATIONAHA.117.027360.

[36] J. Janek and W. G. Zeier, “Challenges in speeding up solid-state battery development,” Nat. Energy, vol. 8, no. 3, pp. 230–240, Mar. 2023, doi: 10.1038/s41560-023-01208-9.

[37] J. Valdovinos, J. Park, J. Smith, and P. Bonde, “Progress on Wireless LVAD and Energy Sources for Mechanical Circulatory Systems,” in Mechanical Support for Heart FailurefJ: Current Solutions and New Technologies, J. H. Karimov, K. Fukamachi, and R. C. Starling, Eds., Cham: Springer International Publishing, 2020, pp. 609–620. doi: 10.1007/978-3-030-47809-4_39.

[38] A. Tsiouris, M. S. Slaughter, A. K. C. Jeyakumar, and A. N. Protos, “Left ventricular assist devices: yesterday, today, and tomorrow,” J. Artif. Organs, Mar. 2024, doi: 10.1007/s10047-024-01436-0.

[39] H.-W. Huang, T.-C. Shih, and C.-T. Liauh, “Predicting effects of blood flow rate and size of vessels in a vasculature on hyperthermia treatments using computer simulation,” Biomed. Eng. OnLine, vol. 9, no. 1, p. 18, Mar. 2010, doi: 10.1186/1475-925X-9-18.

